# Molecular correlates of topiramate and *GRIK1* rs2832407 genotype in pluripotent stem cell-derived neural cultures

**DOI:** 10.1101/710343

**Authors:** Richard Lieberman, Kevin P. Jensen, Kaitlin Clinton, Eric S. Levine, Henry R. Kranzler, Jonathan Covault

## Abstract

There is growing evidence that the anticonvulsant topiramate is efficacious in reducing alcohol consumption. Further, an intronic single nucleotide polymorphism (rs2832407, C ➔ A) in the *GRIK1* gene, which encodes the GluK1 subunit of the excitatory kainate receptor, predicted topiramate’s effectiveness in reducing heavy drinking in a clinical trial. In the current study, we differentiated a total of 22 induced pluripotent stem cell (iPSCs) lines characterized by *GRIK1* rs2832407 genotype (10 A/A and 12 C/C) into forebrain-lineage neural cultures to explore molecular correlates of *GRIK1* genotype that may relate to topiramate’s ability to reduce drinking. Our differentiation protocol yielded mixed neural cultures enriched for glutamatergic neurons. Characterization of the *GRIK1* locus revealed no effect of rs2832407 genotype on *GRIK1* isoform mRNA expression, however a significant difference was observed on *GRIK1* antisense-2, with higher expression in C/C neural cultures. Differential effects of acute exposure to 5 μM topiramate were observed on the frequency of spontaneous synaptic activity in A/A vs. C/C neurons, with a smaller reduction in excitatory event frequency and a greater reduction in inhibitory event frequency observed in C/C donor neurons. This work highlights the use of iPSC technologies to study pharmacogenetic treatment effects in psychiatric disorders and furthers our understanding of the molecular effects of topiramate exposure in human neural cells.

## 1. Introduction

Alcohol use disorder (AUD) is a complex, debilitating, and highly prevalent diagnosis, affecting up to ~14% of the U.S. population in a one-year period (Grant et al., 2015; Grant et al., 2017). The factors contributing to the risk of developing AUD vary among individuals and include both environmental and genetic influences, as well as interactions between the two (Dick and Agrawal, 2008; Gelernter and Kranzler, 2009). Despite the detrimental effects of AUD, the majority of affected individuals never seek treatment (Cohen et al., 2007). Furthermore, the heterogeneity of the disorder among individuals may, in part, contribute to the variable efficacy of currently approved pharmacologic treatments. A better understanding of molecular mechanisms contributing to the heterogeneity of AUD could lead to improved therapeutics and personalized treatments (Heilig et al., 2011).

Topiramate, an anticonvulsant, a prophylactic treatment for migraines, and in combination with phentermine, a weight-loss medication, exerts effects via a broad array of molecular actions that modulate synaptic transmission and neuronal excitability. The medication has also been shown to be efficacious in reducing drinking in individuals with an AUD (Johnson et al., 2003; Johnson et al., 2007; Baltieri et al., 2008; Kranzler et al., 2014). In one study, the ability of topiramate to reduce drinking in treatment-seeking individuals was moderated by a single nucleotide polymorphism (SNP) in the *GRIK1* gene (rs2832407, C ➔ A) (Kranzler et al., 2014). *GRIK1* encodes the GluK1 subunit of the excitatory kainate receptor (Collingridge et al., 2009; Contractor et al., 2011), one of the molecular targets of topiramate (Gryder and Rogawski, 2003; Braga et al., 2009). Kranzler et al. found that topiramate reduced the frequency of heavy drinking days in individuals homozygous for the rs2832407 C allele, while there was no significant difference from placebo among A-allele carriers (Kranzler et al., 2014), an effect that persisted for 6 months following completion of the initial 12-week study (Kranzler et al., 2014). The molecular basis for these long-lasting effects of topiramate in *GRIK1* C-allele homozygotes remains to be elucidated. Because topiramate reduces excitability via its antagonism of GluK1-containing kainate receptors (Gryder and Rogawski, 2003; Braga et al., 2009), the ability of topiramate to reduce drinking in humans may relate to differential effects on synaptic transmission associated with the rs2832407 genotype.

Pluripotent stem cells derived from human fibroblasts (Takahashi et al., 2007) offer the potential to generate multiple cell types of the central nervous system (Mertens et al., 2016) that manifest the donor subject’s genetic background. This method can be used to explore the molecular actions of pharmacogenetic treatments for complex psychiatric disorders in a human cell model system, which may reveal underlying mechanisms leading to differential behavioral effects. Although the application of iPSC technology to study the molecular aspects of personalized medicine for psychiatric conditions is limited, prior work has shown that iPSC-derived neurons can recapitulate differential responses to lithium in bipolar disorder patients, allowing for the exploration of molecular differences underlying the heterogeneity of the treatment response (Mertens et al., 2015; Tobe et al., 2017). Our group has applied iPSC neural differentiation to the study of alcohol use disorder (Lieberman et al., 2012; Lieberman et al., 2015; Lieberman et al., 2017; Jensen et al., 2019) and others have utilized this technology to examine opioid (Sheng et al., 2016) and nicotine (Oni et al., 2016) addiction. However, to our knowledge, no study has utilized iPSCs to advance our understanding of the pharmacogenetic treatment of an addictive disorder.

The objective of the current study was to explore the effects of topiramate in human iPSC-derived neural cultures generated from donors homozygous for the *GRIK1* rs2832407 A or C allele. In total, 10 A/A lines (from 7 donor subjects) and 12 C/C lines (from 10 donor subjects) were utilized to explore genotype-associated differences in expression within the *GRIK1* locus and examine the effects of acute topiramate exposure on synaptic transmission. Results from this study highlight the potential utility of iPSCs for furthering our understanding of pharmacogenetic treatments for complex psychiatric disorders including AUD.

## 2. Experimental procedures

### Generation of iPSCs from GRIK1 rs2832407 A/A and C/C donors

iPSCs were generated from fibroblasts obtained via skin punch biopsies of the inner, upper arm from participants in clinical studies at UCONN Health (UC, Farmington, CT), either non-alcoholic participants enrolled in a study examining the subjective effects of acute alcohol intoxication (Covault et al., 2014; Milivojevic et al., 2014) or from individuals with a DSM-IV diagnosis of alcohol dependence (ADs) enrolled in a pharmacological treatment study examining topiramate’s ability to reduce drinking (Kranzler et al., 2014). Biopsy samples were minced and cultured in Dulbecco’s modified eagles medium (DMEM, Thermo Fisher Scientific) supplemented with 20% fetal bovine serum (FBS, Thermo Fisher Scientific), 1x non-essential amino acids (Thermo Fisher Scientific) and 1x penicillin/streptomycin (Thermo Fisher Scientific). Fibroblast cultures were expanded and passaged using trypsin (Thermo Fisher Scientific) prior to being frozen or sent for reprogramming. Fibroblast DNA was genotyped at rs2832407 using a commercial TaqMan genotyping assay (c_2962029_10, Thermo Fisher Scientific). iPSC lines from 7 A/A and 10 C/C subjects were used for analysis.

The UC Stem Cell Core reprogrammed fibroblasts to pluripotency using retrovirus to express five factors (*OCT4, SOX2, KLF4, c-MYC*, and *LIN28*) or sendai virus to express four factors (*OCT4, SOX2, KLF4*, and *c-MYC*). Two to four weeks after viral transduction, multiple pluripotent stem cell colonies for each subject were selected and expanded as individual iPSC lines. Expression of pluripotency markers by iPSC cells was verified by immunocytochemistry for SSEA-3/4 and NANOG by the UC Stem Cell Core. iPSCs were cultured on a feeder layer of irradiated mouse embryo fibroblasts using human embryonic stem cell media containing DMEM with F12 (DMEM/F12, 1:1 ratio, Thermo Fisher Scientific) supplemented with 20% Knockout Serum Replacer (Thermo Fisher Scientific), 1x non-essential amino acids, 1 mM L-glutamine (Thermo Fisher Scientific), 0.1 mM β-mercaptoethanol (MP Biomedicals), and 4 ng/mL of basic fibroblast growth factor (bFGF, Millipore). Media was fully replaced daily and cells were cultured to confluency before being passaged using 1 mg/mL Dispase (Thermo Fisher Scientific) in DMEM/F12.

### iPSC neural differentiation

iPSCs were differentiated into neural cell cultures as previously described (Lieberman et al., 2012) using a protocol developed by the WiCell Institute for the differentiation of human embryonic stem cells into neural cells of a forebrain lineage (#SOP-CH-207, REV A, www.wicell.org, Madison, WI). We utilized an embryoid body-based protocol wherein iPSC colonies are removed from the feeder layer substrate and cultured in suspension prior to neural induction. Neuroepithelial cells were generated by culturing for 3 weeks in neural induction media containing 1x N2 supplement (Life Technologies) and 2 μg/mL heparin (Sigma Aldrich), following which cells were dissociated and cultured in 24-well plates on poly-L-ornithine and Matrigel (BD Biosciences, Bedford, MA) coated glass coverslips in neural differentiation media containing neural growth factors 1x B27 supplement (Life Technologies), 1 μg/mL laminin (Sigma-Aldrich), and 10 ng/mL each of brain-derived neurotrophic factor (BDNF, Peprotech, Rocky Hill, NJ), glial-derived neurotrophic factor (GDNF, Peprotech), and insulin-like growth factor 1 (IGF-1, Peprotech). All cells were incubated at 37° in 5% CO_2_.

### Immunocytochemistry

Twelve weeks after being plated onto glass coverslips, neural cultures derived from 3 A/A and 5 C/C donors were fixed using 4% paraformaldehyde in PBS for 20 min at room temperature and permeabilized using 0.2% triton X-100 (Sigma-Aldrich) in PBS for 10 min. Following a 1-hr block using 5% donkey serum (Jackson ImmunoResearch, West Grove, PA), cultures were incubated for 24-48 hours at 4° with the following primary antibodies diluted in 5% donkey serum in PBS: mouse anti-beta III-tubulin (1:500, Covance, Dedham, MA), mouse anti-GFAP (1:500, Millipore, Brillerica, CA), rabbit anti-MAP2 (1:500, Millipore), rabbit anti-TBR1 (a forebrain glutamatergic neuromarker; 1:1000, ProteinTech Group, Chicago, IL, incubation included 0.1% triton X-100), and rabbit-anti GluK1 (1:100, Thermo Fischer). Cells were then washed and incubated at room temperature for 2 h in donkey anti-mouse alexa fluor 594 (1:1000, Life Technologies) and donkey anti-rabbit alexa fluor 488 (1:1000, Life Technologies) secondary antibodies diluted in 3% donkey serum in PBS, and mounted in DAPI-containing media for visualization.

### Electrophysiology

Whole cell patch-clamp electrophysiology was performed on neurons differentiated from rs2832407 A/A and C/C iPSCs using previously described techniques (Lemtiri-Chlieh and Levine, 2010; Fink et al., 2017). Neurons were selected for recording based on morphology, including pyramidal-shaped soma and the presence of neurites. Artificial cerebrospinal fluid (aCSF) containing 125 mM NaCl, 2.5 mM KCl, 1.25 mM NaH_2_PO_4_, 1 mM MgCl_2_-6H_2_O, 25 mM NaHCO_3_, 2 mM CaCl_2_, and 25 mM dextrose was perfused through the recording chamber at 1 ml/min at room temperature. An internal recording solution containing 4 mM KCl, 125 mM K-gluconate, 10 mM HEPES, 10 mM phosphocreatine, 1 mM EGTA, 0.2 mM CaCl_2_, 4 mM Na_2_-ATP, and 0.3 mM Na-GTP (pH 7.3) was used for all recordings. Characterization of basal neuronal properties was performed on 15 A/A and 27 C/C neurons derived from 1 A/A and 2 C/C donors and cultured for 17-19 weeks after plating onto glass coverslips. Upon break-in, neurons were noted for their resting membrane potential by injection with 0 current and corrected post hoc for liquid junction potential. Action potentials were evoked in current clamp mode at ~-70 mV by applying 500-ms duration current steps from −20 pA to +40 pA in 5 pA intervals.

The acute effect of topiramate on spontaneous synaptic activity was examined in voltage clamp mode. We explored the effects of 5 μM topiramate in our experiments based on *in vitro* studies showing 50% antagonism of GluK1-containing kainate receptors by 0.5 μM topiramate, consideration of trough serum topiramate concentrations (7-24 μM) in a sample of 344 patients given topiramate 300 mg/d for the treatment of epilepsy (May et al., 2002), and consideration of the experimental design demonstrating moderation by rs2832407 of topiramate’s efficacy to reduce heavy drinking, which utilized a 12-week escalating dose starting at 25 mg/day and increasing weekly to a maximal daily dose of 200 mg (Kranzler et al., 2014). In total, 25 A/A and 30 C/C neurons derived from 3 A/A and 4 C/C subjects and cultured for a minimum of 3 months were used for analysis. Neurons were validated by the presence of voltage-gated inward and outward currents, and ability to fire an action potential. To observe excitatory synaptic activity, neurons were held at −70 mV. To observe inhibitory synaptic activity, neurons were held at 0 mV. A 10-min baseline recording was performed (a 5-min equilibration period and a 5-min experimental period), followed by a 30-min perfusion of aCSF supplemented with 5 μM topiramate. Only one neuron was examined per coverslip following perfusion with topiramate. The last 5 min of the baseline recording period was used for percent-change normalization, and the frequency of synaptic events recorded during topiramate exposure was binned into 10-min intervals for statistical analysis and graphical presentation. Cells were excluded from the analysis if they were unable to fire an action potential or had 30 or fewer spontaneous events during the baseline recording (0.1 Hz). All electrophysiological recordings were performed using a HEKA EPC9 amplifier and PatchMaster software (version 2×67). Analysis and quantification were performed using Axon Clampfit software (version 10.3.1.4).

### RNA extraction and quantitative polymerase chain reaction (qPCR)

Three-month-old neural cultures were used to examine *GRIK1* mRNA expression. In total, 21 neural cell lines derived from 16 donor subjects (10 lines from 7 A/A donors and 11 lines from 9 C/C donors) were used for qPCR gene expression analysis, with RNA from 6 coverslips processed separately per subject. RNA was extracted using TRIzol reagent (Thermo Fisher Scientific) following the manufacturer’s instructions. RNA was quantified using a NanoDrop 2000 spectrophotometer (Thermo Fischer Scientific, Pittsburgh, PA) and cDNA was synthesized from 2 μg RNA using a High Capacity cDNA Reverse Transcription kit (Thermo Fisher Scientific).

cDNA was analyzed by quantitative real-time polymerase chain reaction using an Applied Biosystems 7500 instrument (Thermo Fisher Scientific) and TaqMan Assays-on-Demand (Thermo Fisher Scientific) FAM-labeled probe and primer sets for *GRIK1* isoform 1 (exons 9-10, Hs01081331_m1), *GRIK1* isoform 2 (exons 8-10, Hs01081334_m1 and exons 16-18, Hs01081332_m1), total *GRIK1* (exons 1-2, Hs00168165_m1) and *GRIK1* antisense-2 (Hs00370612_m1). Expression was quantified relative to a VIC-labeled TaqMan probe for the reference gene *GUSB* (4326320E). cDNA synthesized from RNA extracted from each culture well was assayed in triplicate 20-μL reactions containing *GRIK1* FAM-labeled and *GUSB* VIC-labeled assays using Gene Expression Master Mix (Thermo Fisher Scientific) per the manufacturer’s protocol.

PCR cycles were as follows: 95°C for 10 min, followed by 40 cycles of 95°C for 15 sec and 60°C for 60 sec. A standard curve consisting of a 5-level serial dilution of 200%, 100%, 50%, 25%, and 12.5% of a cDNA pool from untreated 12-week old neural cultures differentiated from four subjects was included on each plate to determine the relative mRNA expression across different qPCR plates. Data are displayed as mRNA abundance relative to this cDNA pool and to the housekeeping gene *GUSB* (where a unit of 1 is equivalent to the abundance of the target gene relative to *GUSB* in the reference RNA sample). For each cell line, the expression of specific *GRIK1* isoforms and antisense-2 was normalized to the expression of total *GRIK1*.

### Statistical analysis

Statistical analysis was performed using GraphPad Prism software (V5.0f for Mac, GraphPad Software, www.graphpad.com) or SPSS (v21, IBM, Armonk, NY). Student’s t-tests were used to compare TBR1+ staining, basal neuronal properties, and mRNA expression between A/A and C/C neural cell cultures. Chi-square analysis was used to compare the action potential phenotype between A/A and C/C neurons. Generalized linear mixed models containing *GRIK1* rs2832407 genotype (A/A vs. C/C), topiramate treatment time, and their interaction as factors were utilized to examine the effect acute of topiramate exposure on the frequency of spontaneous synaptic events. Statistical significance was defined as p < 0.05.

## 3. Results

### iPSCs from rs2832407A/A and C/C donors differentiate into functional neurons

Using immunofluorescence, GluK1 could be visualized on the soma and on protrusions residing along neurites of beta III-tubulin-positive neurons (Figure 1A-B) generated from *GRIK1* rs2832407 A/A and C/C donors. The differentiation protocol produced mixed neural cultures containing both MAP2-positve neurons and GFAP-positive astrocytes (Figure 1C). There was no significant difference in the efficacy of *GRIK1* A/A or C/C iPSCs to differentiate into neuronal cultures enriched for TBR1-positive glutamate neurons (p = 0.54) (Figure 1D-E), with ~50% of the total cells staining positive for TBR1.

**Figure 1.**
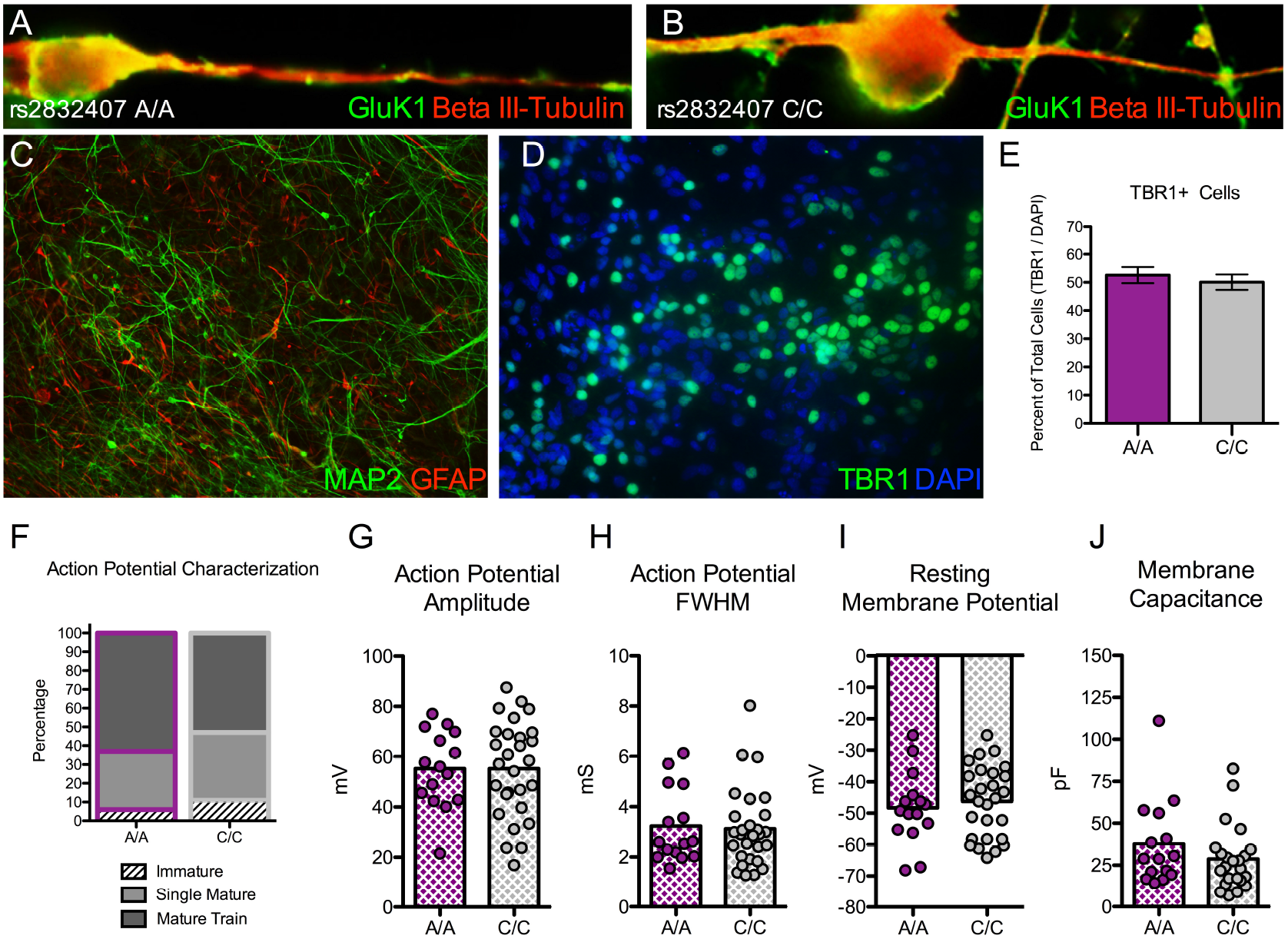
*GRIK1* rs2832407 A/A and C/C iPSCs differentiate into functional neurons. (A-B) A/A and C/C neurons 12 weeks after plating express GluK1 (green), which can be observed localizing to the soma and along beta-III tubulin positive neurites. (C) Neural differentiation produces mixed cultures containing MAP2-positive neurons and GFAP-positive astrocytes. (D-E) Cultures are enriched for TBR1-positive glutamate neurons. The percentage of TBR1-positive cells relative to the total cell population does not differ in neural cultures derived from 3 A/A and 5 C/C donors. 7,163 total cells total were counted for analysis. (F) iPSC-derived neurons (17-19 weeks post plating) generate mature action potential trains in response to a depolarizing current injection. The percentage of neurons that generate immature, single mature, or mature action potentials did not differ between 1 A/A and 2 C/C donors. (G-J) No difference between *GRIK1* genotypes was observed in neuronal properties including (G) action potential amplitude, (H) action potential full width at half-maximum (FWHM), (I) resting membrane potential, and (J) membrane capacitance. Each dot on the graphs depicting distribution represents an individual neuron.

Neurons from A/A and C/C donors were characterized using whole-cell patch clamp electrophysiology. Upon the injection of a depolarizing current, iPSC-derived neurons generated an action potential, with the majority of neurons from both donor groups exhibiting mature action potentials. The percentage of neurons generating an immature, single mature spike, or a mature train did not differ between neurons from the A/A and C/C donors examined (A/A; mature train: 63%, single mature: 32%, immature: 6%, C/C; mature train: 53%, single mature: 36%, immature: 11%, χ^2^ = 0.35, df = 2, p = 0.8) (Figure 1G), and there were no significant differences in either action potential amplitude (t = 0.1, df = 40, p = 0.93) (Figure 1H), or action potential full width at half-maximum (t = 0.1, df = 40, p = 0.93) (Figure 1I). We also did not observe differences between genotypes in resting membrane potential (t = 0.55, df = 40, p = 0.59) (Figure 1J) or membrane capacitance (t = 1.42, df = 40, p = 0.16) (Figure 1K). Taken together, the immunostaining and electrophysiology results showed that our iPSC differentiation protocol was able to generate functional neural cultures from both A/A and C/C donors.

### Gene expression characterization of the GRIK1 locus

Alternative splicing of the *GRIK1* transcript gives rise to GluK1-containing kainate receptors with different C-terminal domains and conductance properties (Hirbec et al., 2003; Jaskolski et al., 2005). We used TaqMan probe and primer sets spanning specific exon/intron boundaries to investigate the total *GRIK1* and isoform-specific expression in 12-week old neural cells differentiated from 10 A/A and 11 C/C lines. We also examined the expression of an antisense RNA (*GRIK1-AS2*) that resides ~800 bp from rs2832407 (Figure 2A). rs2832407 genotype did not associate with differences in mRNA expression of *GRIK1* isoform 1 (exons 9-10 probe: t = 0.1, df =19, p = 0.92), isoform 2 (exons 8-10 probe: t = 0.1, df = 19, p = 0.92; exons 16-18 probe: t = 0.26, df = 19, p = 0.80), or total *GRIK1* (exons 1-2 probe: t = 0.21, df = 19, p = 0.84) (Figure 2B-E). There was a significant difference in expression of *GRIK1* antisense-2 as a function of rs2832407 genotype (t = 2.13.9, df = 19, p = 0.048), with higher expression observed in neural cultures generated from C/C lines (Figure 2F)

**Figure 2.**
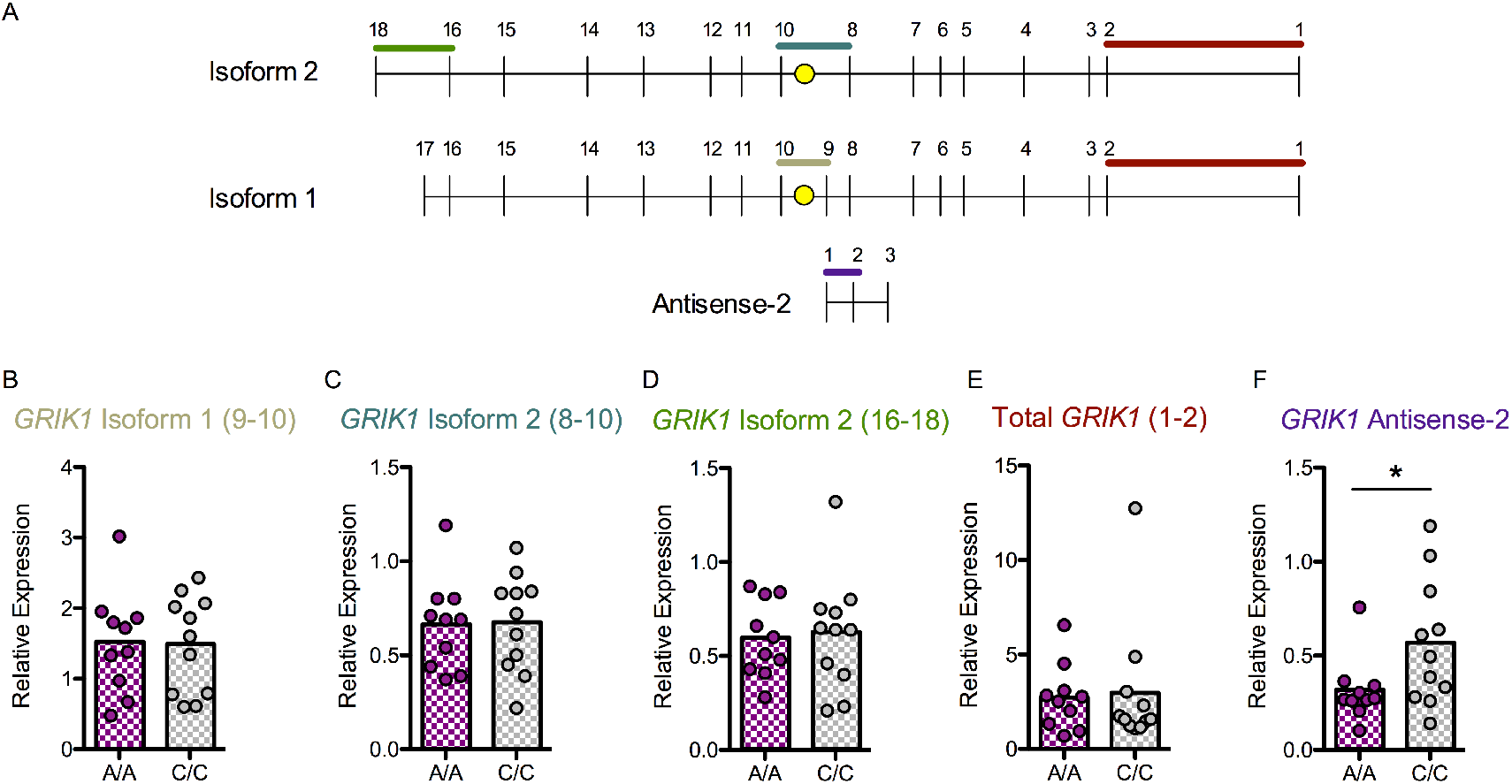
*GRIK1* gene expression in A/A and C/C iPSC-derived neural cultures. (A) Schematic representing the two major *GRIK1* isoforms, which differ by the inclusion of exons 9, 17, and 18, and the antisense-2 RNA. Approximate location of rs2832407 is indicated by the yellow dot, which resides in the intronic region between exons 9-10 in isoform 1 and 8-10 in isoform 2. (B-F) iPSC-derived neural cultures were examined 12 weeks post plating. No significant effect of rs2832407 genotype was observed on the expression of *GRIK1* isoform 1 (B), isoform 2 (C, D), or total *GRIK1* (E). A significant effect of genotype was observed on the expression of *GRIK1* antisense-2 (F). Each dot represents the averaged RNA expression level per cell line.

### Effects of acute topiramate exposure on frequency of synaptic events

GluK1-containing receptors have been reported to enhance the release of neurotransmitter from both excitatory (Aroniadou-Anderjaska et al., 2012) and inhibitory (Braga et al., 2003) presynaptic terminals. To examine genotype effects on presynaptic release in the cultures, we examined the frequency of spontaneous excitatory and inhibitory events following bath application of topiramate. Whole cell patch-clamp recordings of spontaneous excitatory and inhibitory postsynaptic currents (EPSCs and IPSCs, respectively) were obtained from neurons differentiated from 3 A/A and 4 C/C cell lines. EPSCs were recorded from 19 A/A (3 subjects) and 20 C/C (4 subjects) neurons at a holding potential of −70 mV. EPSCs could be blocked by the application of the AMPA/kainate receptor antagonist DNQX (Figure 3A). A significant interaction of genotype with time was observed for EPSC frequency following the perfusion of 5 μM topiramate (F = 5.1, df = 1, 152; p = 0.026), with C/C neurons showing a smaller reduction in EPSC frequency than A/A neurons (~5% vs. ~30% reduction, respectively, after a 30-min exposure to topiramate) (Figure 3B).

**Figure 3.**
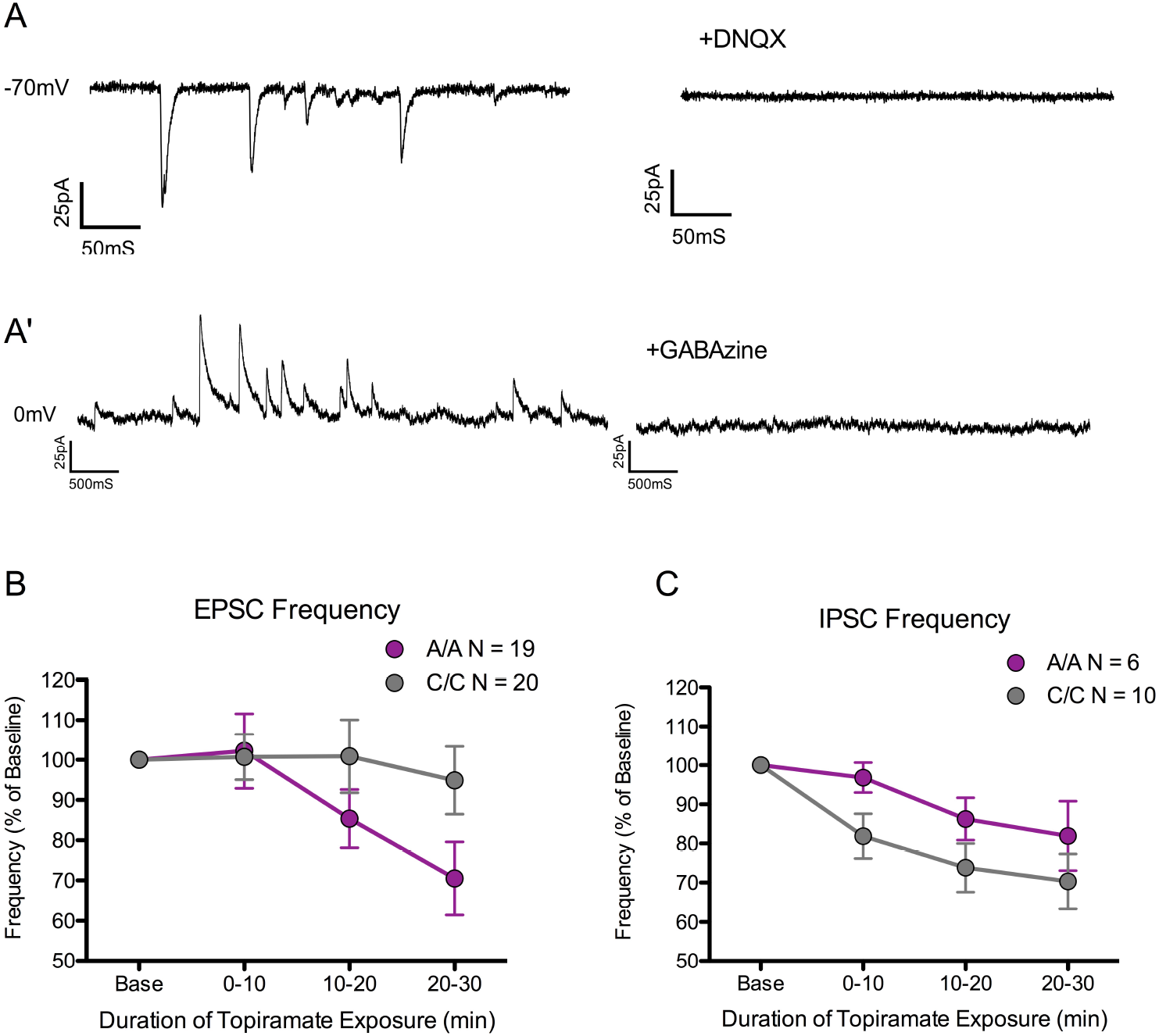
Topiramate exposure attenuates the frequency of spontaneous synaptic activity. (A) Example trace of spontaneous excitatory postsynaptic currents (EPSCs) that are blocked by the AMPA/kainate receptor antagonist DNQX (10 μM). (A’) Example trace of spontaneous inhibitory postsynaptic currents (IPSCs) that are blocked by the GABA_A_ receptor antagonist GABAzine (5 μM). (B-C) Neurons generated from A/A and C/C donors were exposed to 5 μM topiramate for 30 min and spontaneous currents were measured. There were significant interactive effects of topiramate exposure time and *GRIK1* genotype on the frequency of EPSCs and IPSCs.

IPSCs were recorded from 6 A/A (2 subjects) and 10 C/C (2 subjects) neurons at a holding potential of 0 mV. IPSCs could be blocked by the application of the GABA_A_ receptor antagonist GABAzine (Figure 3A’). A significant interaction of genotype with time (F = 5.0, df = 1, 60; p = 0.030) was also observed for the frequency of IPSCs (Figure 3C), with C/C neurons showing a greater decrease in IPSC frequency than A/A neurons in response to topiramate.

## 4. Discussion

Kainate receptors are a class of excitatory glutamate receptors comprised of tetrameric combinations of five subunit types (GluK1-5), with complex biological roles related to temporal changes in regional expression and involvement in both postsynaptic and presynaptic neurotransmission (Contractor et al., 2011). Topiramate, initially approved by the FDA as an anticonvulsant, has been found to reduce alcohol consumption in treatment-seeking individuals (Johnson et al., 2003; Johnson et al., 2007; Kranzler et al., 2014) and non-treatment-seeking volunteers (Miranda et al., 2008). The pharmacologic actions of topiramate are diverse, with *in vitro* studies highlighting inhibitory effects on voltage-gated sodium channels (Zona et al., 1997), L-type calcium channels (Zhang et al., 2000), and non-NMDA glutamate (AMPA/kainate) receptors (Gibbs et al., 2000; Braga et al., 2009), and potentiating effects on GABA_A_ receptors (White et al., 2000; Braga et al., 2009). The inhibitory effects of topiramate on non-NMDA glutamate receptors appear to be mediated primarily by kainate receptors containing the GluK1 subunit (Gryder and Rogawski, 2003; Braga et al., 2009), encoded by the *GRIK1* gene. A single nucleotide polymorphism (rs2832407, C ➔ A) located in intron 9 of *GRIK1* was associated with the risk of alcohol dependence in European Americans (Kranzler et al., 2009) and moderated the efficacy of topiramate in reducing heavy drinking days in treatment-seeking European Americans with AUD (Kranzler et al., 2014; Kranzler et al., 2014). However, the biological mechanisms underlying rs2832407’s contribution to the risk of psychopathology and topiramate’s efficacy in reducing alcohol consumption are unknown.

In the current study, we utilized iPSCs to identify molecular correlates of the *GRIK1* SNP in human neural cell cultures. The *GRIK1* gene (21q22.1) is comprised of 18 exons, generating multiple mRNA isoforms and two antisense RNAs. The two most common isoforms in humans differ by the exclusion of exons 9 and 17 in isoform 2, while isoform 1 contains exons 1-17 but lacks exon 18 (Barbon and Barlati, 2000). Investigation in rodents revealed that alternative splicing of exon 17 and 18 results in amino acid sequence differences in the C-terminal domain, influencing membrane trafficking of the receptor (Jaskolski et al., 2005). Furthermore, the inclusion of exon 18, but not 17, results in PDZ-binding and PKC-phosphorylation sites that can alter kainate receptor localization and function at the synapse (Hirbec et al., 2003). Because of the molecular consequences of alternative splicing, we investigated whether rs2832407 genotype associated with differential *GRIK1* isoform expression, but found no difference in mRNA expression of isoform 1, isoform 2, or total *GRIK1* between A/A and C/C neural cultures.

We did observe a significant association of *GRIK1* genotype and antisense-2 expression, with higher expression in neural cultures derived from C/C donors. Antisense RNAs can influence the expression of genes and proteins via complementary binding to target RNAs, resulting in RNA degradation and/or the inhibition of translation (Saberi et al., 2016). Because antisense-2 RNA was elevated in neural cultures derived from C/C donors, an intriguing possibility is that rs2832407, or functional SNPs in linkage disequilibrium with it, alter the availability of GluK1-containing kainate receptors in C/C subjects by affecting antisense RNA expression, thereby altering the effects of topiramate. The effects of *GRIK1* genotype on antisense-2 expression specifically and GluK1 subunit availability was previously proposed by Kranzler et al. (2014a) as a potential mechanism underlying the pharmacogenetic effects of topiramate on the reduction of alcohol consumption given the proximity of the rs2832407 haplotype block and *GRIK1* antisense-2 transcription start site, including being in near complete linkage with rs363431 that is located within the antisense transcript.

Acute exposure to 5 μM topiramate significantly reduced the frequency of EPSCs and IPSCs. The time course of topiramate effects in reducing EPSC and IPSC frequency is consistent with the reported 10-20 minute delay following bath application of topiramate for the inhibition of kainate receptors (Gibbs et al., 2000; Braga et al., 2009). This slow onset of inhibition is thought to relate to topiramate’s molecular mechanism of action via binding to kainate receptor phosphorylation site(s) when in the dephosphorylated state (Gibbs et al., 2000; Angehagen et al., 2004).

Interestingly, we also observed significant interactions between topiramate treatment and *GRIK1* genotype on the frequency of EPSCs and IPSCs, with an opposite genotype relationship for EPSCs and IPSCs. In response to acute topiramate exposure, neurons derived from C/C donors showed a smaller reduction in EPSC frequency than A/A donors, while neurons from C/C donors had a somewhat greater decrease in IPSC frequency than A/A neurons. The effect of topiramate in reducing EPSCs is consistent with reports that GluK1-containing receptors on presynaptic terminals enhance glutamate release and their inhibition decreases principal neuron EPSC frequency (Aroniadou-Anderjaska et al., 2012). Our results suggest that basal GluK1 facilitation of glutamate release from excitatory presynaptic nerve terminals is reduced in C/C compared with A/A cultures. Similarly, GluK1-containing receptors on GABAergic presynaptic terminals of inhibitory interneurons enhance GABA release at basal levels of extracellular glutamate (Braga et al., 2003), with our results suggesting that this enhancement is less affected by genotype than for glutamate terminals but that there is greater basal facilitation in C/C cultures. With respect to our gene expression findings, it may be that *GRIK1* rs2832407 genotype effects on antisense-2 expression and GluK1 availability would be more pronounced at presynaptic excitatory synapses because we observed a larger reduction in EPSC frequency in neurons from A/A than C/C subjects following topiramate perfusion, suggesting a limited effect of topiramate on presynaptic release at excitatory synapses in C/C neurons. As topiramate’s anticonvulsant effects occur via modulation of the balance between excitatory and inhibitory neurotransmission, it is interesting to speculate that rs2832407 genotype effects related to the greater reduction in heavy alcohol consumption in C/C subjects by topiramate (Kranzler et al., 2014; Kranzler et al., 2014) may be due, in part, to differential presynaptic effects of topiramate as a function of rs2832407 genotype. Alternatively, *GRIK1* genotype and consequent biological functions may have developmental effects on the relative strength of interactions between brain regions that contribute to both an association of genotype with alcohol dependence and the reduction in drinking produced by topiramate.

The findings reported here must be viewed in the context of several limitations. First, we examined the effect of acute topiramate exposure on non-pharmacologically isolated spontaneous synaptic events that contained a mixture of action potential-dependent and non-action potential-dependent events. Therefore, it may be that the synaptic effects of topiramate observed in our iPSC-derived neurons were due to topiramate’s effect on receptors or channels other than GluK1-containing kainate receptors, e.g., voltage-gated sodium and calcium channels residing on the non-voltage clamped presynaptic neurons, although such components would not be expected to show *GRIK1* genotype effects. Methods developed in *in vitro* rodent models to pharmacologically isolate kainate receptors (Gryder and Rogawski, 2003; Braga et al., 2009) may be more useful for examining the specific effects of topiramate in iPSC-derived neurons. Second, because iPSC neural cultures more closely resemble early brain development than adult neural tissue, with transcription profiles from iPSC neural cultures most closely resembling those of first-trimester brain tissue (Brennand et al., 2015), the effects of topiramate and *GRIK1* genotype identified in this culture system may not reflect their effects in mature neural tissue. The neural differentiation protocol we utilized generates cultures enriched for forebrain-type glutamate neurons. Because the expression of kainate receptors varies by brain region (Contractor et al., 2011) and the effects of topiramate differ between excitatory and inhibitory neurons (Braga et al., 2009), future work should consider use of additional neural differentiation protocols to generate cultures enriched for inhibitory, excitatory, or dopaminergic neurons, among others (Mertens et al., 2016), to explore the effects of topiramate treatment on other neuronal cell types. The potential value of examining neurons from additional differentiation protocols is highlighted by our prior work (using the same forebrain lineage cell differentiation protocol) (Jensen et al., 2019) in which we observed that previously identified eQTLs from human cortical tissue had greater eQTL effects within the neural cultures than previously identified eQTLs from other subcortical brain regions and peripheral tissue. Finally, it remains to be determined how the effects of topiramate observed in our culture system in A/A and C/C donors relate to the differential efficacy by genotype in reducing heavy drinking. We expect that iPSC technologies will also be a valuable tool for examining the molecular mechanisms underlying the pharmacogenetic effects of other pharmacological treatments for AUD (Jones et al., 2015).

In summary, we have shown that iPSC-derived neural cells are a model system to explore the molecular actions of topiramate in relevant cell types *in vitro*. In particular, the technology allows one to probe mechanisms of pharmacogenetic treatment effects, even though the functional genetic element is not yet known, by utilizing cells that express the unmodified donor genome. Future studies can use novel protocols to generate specific populations of neural derivatives to identify between-cell pharmacologic sensitivity and the influence of genetic variation on neural activity.

## Acknowledgments

We would like to thank Leann Crandall at the UC Stem Cell Core for her valued assistance in generating iPSC lines.

